# Multicore-fiber microendoscopy for functional cellular in-organ imaging

**DOI:** 10.1101/2024.03.02.583077

**Authors:** Tobias A. Dancker, Mohamed Ibrahem Elhawy, Ramona Rittershauß, Qinghai Tian, Yvonne Schwarz, Markus D. A. Hoffmann, Christopher Carlein, Amanda Wyatt, Vanessa Wahl, Daniel Speyerer, Alaa Kandah, Ulrich Boehm, Leticia Prates Roma, Dieter Bruns, Peter Lipp, Gabriela Krasteva-Christ, Marcel A. Lauterbach

## Abstract

Microendoscopy enables minimally invasive investigations of organs even within small cavities. Conventional microendoscopy is limited by probe size and often restricted to a single excitation wavelength. We developed and characterized a multichannel microendoscope as thin as 360 µm and recorded functional cellular signals in-situ using custom written software for image processing. The endoscope had an effective resolution of 4.64 µm and resolved subcellular structures of neurons. The system enabled analysis of in-situ calcium responses in murine tracheal brush cells and kidney podocytes. Additionally, ratiometric redox responses were recorded in whole, explanted organs and pancreatic islet culture. The flexibility and simplicity of our approach for imaging a variety of tissues and organs paves the way for in-vivo, longitudinal studies with cellular resolution.

## 1. Introduction

### 1.1 Endoscopy

Endoscopy is an established method for minimally invasive imaging in humans and animals and is often employed concomitant to the collection of biopsies [1, 2]. If an endoscope is equipped with suitable light sources and filters, it can also be used to examine the fluorescence characteristics of tissues. This can help in the stratification of tumors, either through their autofluorescence spectrum or following probing with specific fluorescent dyes [1, 3]. Given the ready availability of many genetically modified mouse lines expressing fluorescent proteins and the unlimited possibilities for gene manipulation in mouse models, the application of endoscopy in mice is highly attractive. However, for the application in preclinical rodent models, the use of available endoscopes is limited due to their probe size and thus microendoscopes would be more suitable. Recently described microendoscopes are limited by a fiber diameter exceeding 500 µm, which prohibits the application in small organs or lumina of mice [4–8]. A microendoscope of suitable size with multiple channels and with the possibility to switch illumination wavelengths would be of great value and allow the minimally invasive examination of tissues and/or organs, enabling longitudinal studies with reduced sample sizes. These studies may include – but will not be restricted to – the investigation of cellular or molecular responses to drugs, cell metabolism, or the response to environmental factors.

Systems whose diameter is compatible with mouse models have been published as early as 2013 but often lack multiple excitations and flexibility [9–14]. While a size-compatible system with a single excitation source is useful, it limits the microendoscope to a single-channel imaging. With continuously evolving organic fluorophores and fluorescent proteins this is a limiting factor.

Several published systems are capable of imaging cells [6, 9–12, 15, 16] and some are even capable of multimodal imaging, multichannel imaging, or present impressive advances in lens technology. However, these systems have either never been used to image animals or are too large to consider applying their technological advantages in the smaller organs or lumina of mice.

The microendoscope introduced in this study is equipped with multiple excitation wavelengths, has a maximum diameter of 360 µm and provides cellular resolution at a working distance of 110 µm. Employing this microendoscope, we recorded functional signals such as calcium signals in brush cells of the intact trachea and in podocytes of the intact kidney, from whole mice in-situ. In addition, mitochondria-associated redox signals were recorded in the excised pancreas and kidney. Furthermore, redox signals were ratiometrically analyzed in isolated and cultured pancreatic islets. Neurons expressing a green calcium indicator and a red fluorescent protein were used to exploit the ability of the microendoscope to image with a reference channel.

### 1.2 Tracheal Brush cells

Tracheal brush cells were only recently recognized as chemosensory cells in the trachea [17–19]. They owe their name to brush-like villi that extend into the tracheal lumen and have been shown to detect bitter tasting substances in the trachea and regulate respiratory rate via cholinergic transmission [20]. These signals act as paracrine messengers, with autocrine regulation [21]. Brush cell activation has also been shown to induce a local innate immune response such as recruitment of neutrophils from the peripheral blood via stimulation of nociceptor sensory neurons, suggesting its relevance as a defense mechanism against infection [22]. In the same study, denatonium benzoate was found to be an agonist of brush cells, triggering a transduction cascade via the transient receptor potential cation channel subfamily M member 5 (TRPM5), which causes an increase in intracellular calcium concentration. Due to its small diameter, our microendoscope is uniquely equipped for imaging these calcium signals directly in GCaMP-expressing brush cells in the whole mouse.

### 1.3 Podocytes

Podocytes are part of the glomeruli and are responsible for the primary filtration of blood in the kidney. The foot processes of the podocytes form the filtration slits, whose selective permeability ensures that no cellular component leaves the blood [23]. Old or injured podocytes lose their density and shape, allowing more material to leak from the blood into the primary urine [24, 25]. Calcium signaling cascades in podocytes are responsible for regulating the number of foot processes, their density, structure, and cytoskeleton. These signals are regulated by angiotensin 2 stimulation, with injured podocytes becoming more sensitive [26–28]. Rather than isolating the podocytes or preparing kidney slices for conventional microscopy the microendoscope is capable of imaging GCaMP3 signals in the intact kidney without further mechanical manipulation.

### 1.4 Mito-roGFP2-Orp1

Reduction/oxidation-sensitive green fluorescent protein 2 (roGFP2) is a ratiometric fluorophore based on the jellyfish-derived green fluorescent protein (GFP), modified to exhibit different excitation properties depending on the molecule’s oxidation status [29]. Depending on the state of the fluorophore it is more efficiently excited by ultraviolet (UV) or by blue light. Due to this property, roGFP2 has proven to be a valuable tool for the ratiometric investigation of the redox dynamics in several cellular models [30, 31]. Through genetic targeting, the roGFP2-Orp1 sensor can be expressed in various cellular compartments such as the cytosol, mitochondrial intermembrane space or mitochondrial matrix (mito-roGFP2) [32]. Additionally, through fusion with the peroxidase-1 protein (Orp1) the combined molecule (mito-roGFP2-Orp1) can be used to image H_2_O_2_-specific fluorescence changes [33]. This biosensor can be used to study redox changes in isolated structures or whole tissues. The microendoscope can illuminate a cell or organ expressing mito-roGFP2-Orp1 with 395 nm light in one frame and 475 nm light in the next, allowing for ratiometric imaging.

Applying the microendoscope for functional imaging in the kidney, trachea and pancreas, we demonstrate the ability of the instrument to reliably image quantifiable signals of individual cells in single and multi-channel acquisitions in several differently challenging mouse organs and models.

## 2. Methods

### 2.1 Microendoscope hardware and software

We designed our microendoscope [Fig. 1(a–d)] with the aim to achieve a minimal probe diameter (360 µm) while still being able to image at a working distance that is sufficient for deep cellular imaging within the examined organs. This is achieved by using a single gradient index (GRIN) lens fused to the end of a multicore fiber [Fig. 1(d)]. A mechanical three-axis micromanipulator served for positioning the imaging end of the fiber with the GRIN lens above the sample. A syringe cannula (inner diameter 0.60 mm), the ends of which had been filed off, was used as a guide tube [Fig. 1(d)]. To allow for larger movement in animal experiments and to reach otherwise inaccessible crevices of e.g., the trachea we constructed a tilting sample/animal stage using two posts and a 90° post clamp. With this stage the specimen can be rotated along the horizontal and vertical axes, and coarsely adjusted in height [Fig. 1(a)].

**Fig. 1.**
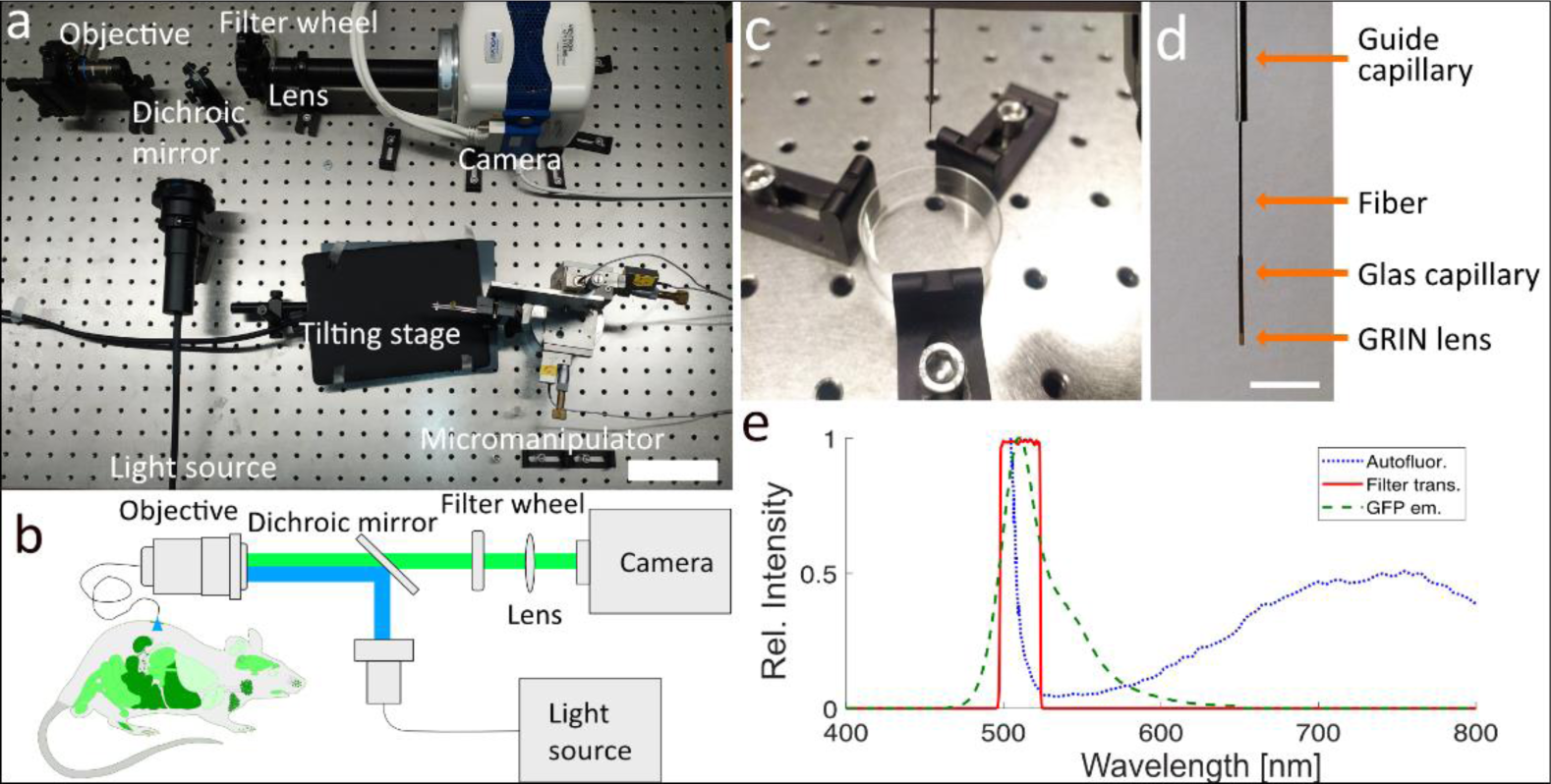
Microendoscope-system. a) Top view of the optical table (breadboard) set-up, in view are the camera, light guide coming from the light source, the micromanipulator, the tilting stage and the optical components. Scale bar 10 cm. b) Light path of the microendoscope-system. c) Fiber end in the micromanipulator over a cell culture dish. d) Close up of the fiber end, in view are the GRIN lens at the end of the fiber, the glass capillary holding it in place and the syringe cannula serving as guide. Scale bar 5 mm. e) Fiber autofluorescence spectrum [34] (dotted blue) overlaid with the emission spectra of GFP (stippled green) and the transmission of the 511/20 nm emission filter (red).

We assembled the microendoscope on a 600 x 900 mm breadboard (Newport, Darmstadt, Germany) [Fig. 1(a, b)] using standard optomechanical parts obtained from Thorlabs (Thorlabs, Newton, New Jersey, United States). The system uses a Fujikura Imaging fiber bundle (FIGH016-160S) with a nominal 145 μm image circle, 1600 ± 10% fibers and 210 μm coating diameter. Attached to one end is an adapted GRIN lens (NEM-025-06-00-520-S), with 250 µm diameter, and 110 μm working distance, an NA of 0.5 and a nominal magnification of −1. These parts were fixed with optical adhesive and secured with a polyimide-coated glass capillary 360 µm in diameter and 7 mm in length by Grintech (Jena, Germany) [Fig. 1(c, d)]. This capillary defines the thickest part of the endoscope.

The other end of the fiber bundle is imaged in a 4f configuration onto a CCD camera, using a 50x objective with a focal length of 3.6 mm (MPLFLN50X, Olympus, Shinjuku City, Japan, no immersion, NA 0.8) and a doublet lens with a focal length of 200 mm. The fiber was mounted in a 6-axis kinematic mount (Thorlabs) [Fig. 1(a, b)].

For fluorescence imaging using the Fujikura Imaging fiber bundle, its autofluorescence spectrum must be considered, especially since it spans the spectral range of common fluorescent proteins. We used the results of Udovich and coworkers who measured the autofluorescence spectrum with 488 nm excitation [34] to optimize filters and the dichroic mirror. The autofluorescence spectrum shows two main peaks, one in the blue-green spectral region below 504 nm and one at 754 nm [Fig. 1(e)] [34]. On the one hand, we wanted the option to flexibly image multiple emission bands and fluorophores without switching filters between channels (see below). On the other hand, many of our mouse lines are based on GFP variants without additional fluorophores, which allows to exclude the long-wavelength peak of the autofluorescence. We thus opted for a design with a fixed multi-band dichroic (F67-401, AHF, Tübingen-Pfrondorf, Germany, Supplementary Fig. S1), facilitating stable alignment, but two alternative emission filters. For multi-emission imaging, we used a 435/15, 520/10, 595/15, 695/30 quad band emission filter (F67-401, AHF). For imaging of GFP-variants (e.g. GCaMP, roGFP2) in the absence of other fluorophores we used a 511/20 single band filter (F39-509, AHF) which rejected the red autofluorescence peak [Fig. 1(e)].

The light source was a Lumencor Aura Lightengine (Beaverton, Orlando, USA) with LEDs and fixed excitation filters at 395/25, 475/28, 555/28, 635/22 and 730/40 nm. The camera was an Evolve 512, monochrome, 16-bit, CCD camera (Photometrics, Tucson, AZ, USA).

Electronics of the microendoscope were constructed to allow for frame-by-frame alternating excitation (frame-interleaved) with different wavelengths. This allowed us to reliably measure multiple color channels without any spatial shift between channels and without any (slow) switching of filters: To control the excitation light source an Arduino UNO (www.arduino.cc) was programmed to set which excitation light source is active during which frame. The Arduino specific programming language was used. During every frame, the camera generates a frame exposure TTL signal. The Arduino computer splits these TTL signals to the different excitation LEDs in the Lightengine. Which LED activates during which frame can be pre-configured by the user via a graphical user interface (GUI). Sequential excitations can be configured within this GUI for framewise excitation switching. When a time series is completed and no TTL signal from the camera is detected for 5 s, the excitation sequence is automatically reset. This ensures a consistent assignment of the color channels by the Arduino so that each recording starts with the same excitation wavelength.

Micromanager 2.0 was used for camera control and image acquisition. Image processing was performed with custom-written routines in MATLAB (Mathworks, Natick, USA).

### 2.2 Image processing

To optimize image analysis beyond the raw images, we designed an image processing pipeline to process our recordings (Fig. 2). This pipeline accomplished two main tasks: 1) Removal of the individual fiber cores in the image, and 2) removal of the fiber autofluorescence. In raw recordings the individual cores of the multicore fiber are still visible with their cladding. These spaces between the cores are regions of constant intensity that will skew any intensity analysis of more than one core.

**Fig. 2.**
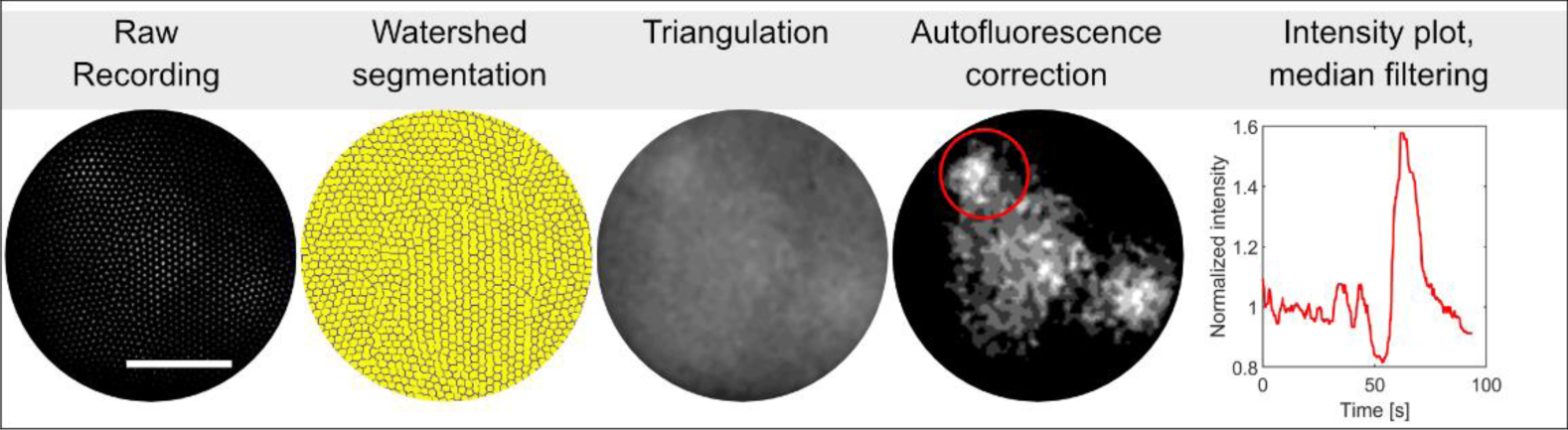
Processing steps of images acquired with the microendoscope. Raw images generated via the multicore fiber are segmented by a watershed function, which identifies the borders of each core. Intensities of all pixels in each core are averaged and a smooth image is calculated by triangulation-based linear interpolation. This image still contains the fiber autofluorescence, which is removed by subtracting the 25^th^ intensity percentile of each frame. Next an ROI-based intensity profile in the time dimension is plotted and filtered with a sliding window median filter with a window size of 30 frames (approximately 6 seconds). Scale bar 50 µm.

To obtain a smooth image from which the honeycomb pattern of the individual fiber cores was removed, the raw images were processed with custom-written MATLAB routines using a watershed segmentation (MATLAB command watershed) to find each individual fiber core. To determine the positions of the fiber cores a reference image in which the fiber cores are well resolved was used (either 395 nm excitation, leading to strong autofluorescence or an image of a homogeneous sample, or the average of a longer time series) (Fig. 2, left panel). The reference image was smoothed by convolution with a two-dimensional Gaussian function. Regions of individual cores were identified after water shedding and segmented. In each region covered by an individual fiber core, the fluorescence intensity was averaged. Triangulation-based linear interpolation was used to construct an interpolated image for each frame of the time series. The image was interpolated to only 128 x 128 pixels to reduce the unnecessary oversampling of the individual cores of the fiber bundle in the resulting image, reducing the amount of data to be stored and handled.

To enhance contrast and reduce noise, we subtracted the 25^th^ percentile intensity value from every frame so that the fluorescence of cells remains comparably bright, while regions of pure autofluorescence are darkened. The dark corners outside the round fiber, which are not part of the field of view, were not included in the percentile calculation. The 25^th^ percentile was chosen, because for the sparse labeling present here, it represents the dark (“signal free”) regions of the image well, but it is less susceptible to noise than for example, the minimum intensity of the images. An individual correction was applied to each image rather than subtracting a common calibration image to prevent camera flicker artefacts from interfering with the detection of cell-specific fluorescence. Since we expected relatively bright autofluorescence, a small instability in the overall fluorescence intensity that is recorded (e.g., unstable detection efficiency, dwell time, camera gain or illumination), multiplied by the large autofluorescence intensity could lead to a flickering artifact and affect the intensity plot, so we chose to subtract a proportion per frame rather than one base value.

To analyze functional fluorescence signals on the processed and autofluorescence corrected images, cell specific regions of interest (ROIs) were manually drawn and intensity profiles of ΔF/F ([spatial mean of intensity within the ROI over the serieś lowest intensity]-1) over time were calculated. This intensity trace was filtered using a moving window median filter with a window size of 30 frames (comparison of raw data/median filtered data shown in Supplementary Fig. S2).

To calculate standard deviation between these signals, alignment to the signal onset was necessary. The onset of the intensity increase was identified by finding the first point in time where intensity increased above 80% of the maximum intensity. All signals were aligned to this time point.

### 2.3 Optical characterization

To visualize and count all fibers of the bundle [Fig. 3(a)], a reference image with 395/25 nm excitation and 511/20 emission filter showing the fiber autofluorescence was acquired without a sample with an exposure time of 200 ms. The watershed segmentation (see above) was used to count the fiber cores.

**Fig. 3.**
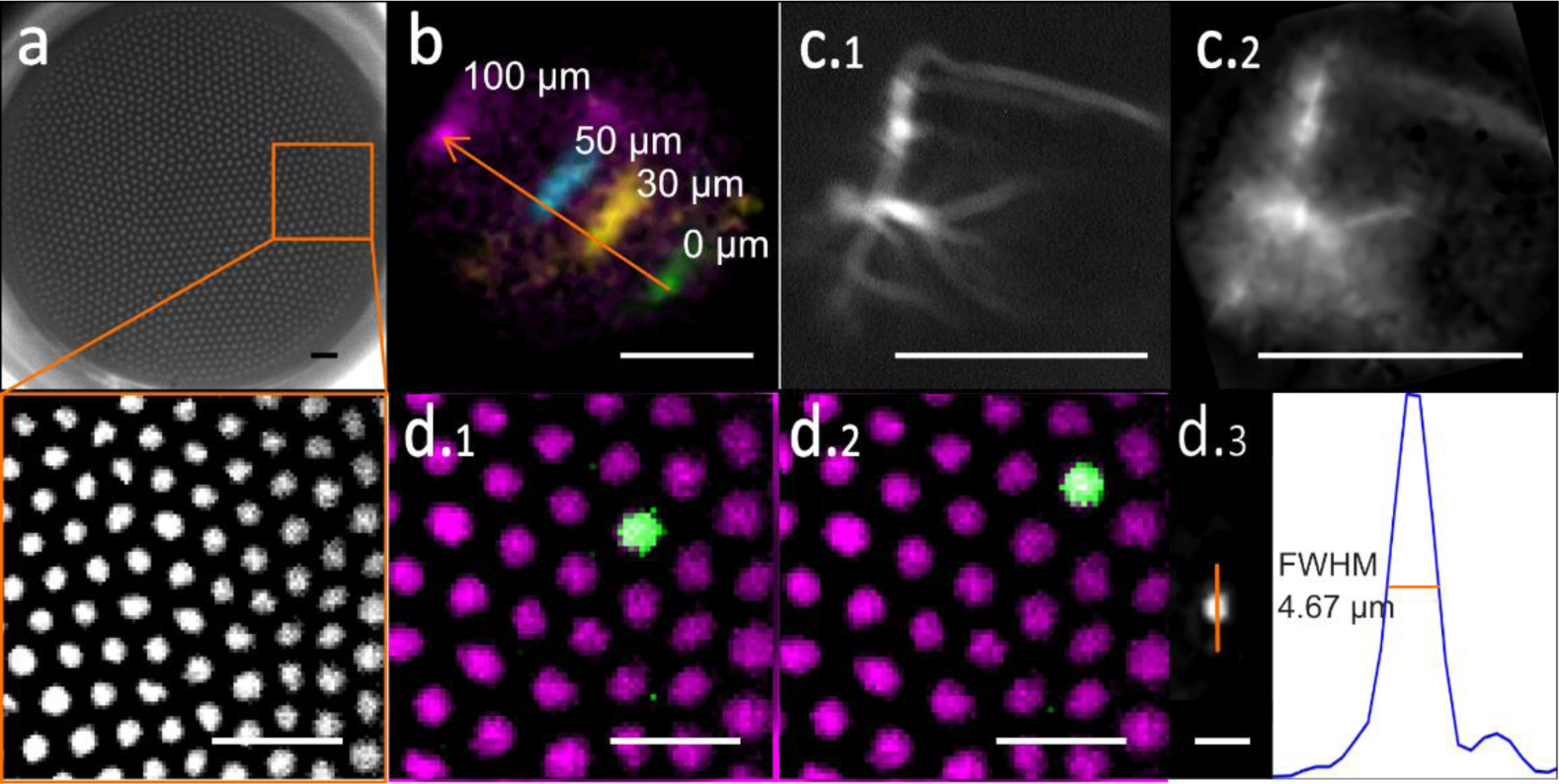
Optical characterization of the microendoscope. a) Image of the microendoscope fiber bundle showing a single fiber core to be 2.3 µm in diameter. Scale bars 10 µm. b) Calibration of magnification and FOV: test sample (fluorescent fibers) before (green) and after a shift of 100 µm (purple), 50 µm (cyan) and 30 µm (yellow) using the micromanipulator. The shift on the camera was used to calculate the magnification and FOV. Scale bar 50 µm. c) Proving single-cell resolution by comparative images of the same neuron taken simultaneously with a conventional microscope (c.1) and the microendoscope (c.2). Scale bars 100 µm. d) Resolution measurement with microspheres: d.1) in an unprocessed image a 1 µm bead (green) appears only on one of all fibers (magenta, intensity-scaled reference image without sample). d.2) The bead becomes visible on the neighboring fiber after shifting by one core distance. d.3) corresponding processed image (honeycomb pattern removed) with intensity profile through the bead. Scale bars in d.1 and d.2 10 µm, in d.3 5 µm.

To characterize the field of view (FOV) and magnification of the microendoscope, fluorescent text marker (Stabilo Boss Pink, Heroldsberg, Germany) was applied onto optical cleaning tissue (MC-5, Thorlabs) and allowed to dry. The fibers of the tissue were imaged with the microendoscope using the 475/25 nm excitation. Using the micromanipulator, the sample was moved by 30 µm, 50 µm and 100 µm [Fig. 3 (b)]. The number of pixels that the fibers moved was calculated using the Pythagorean theorem. The movement of the probe divided by the number of pixels was used to calculate the size of the pixel. To calculate the magnification, the distance in pixels was multiplied by 16 µm (the size of the pixel on the camera chip) and then divided by the distance moved with the micromanipulator. The standard error of the three measurements was calculated.

To compare the microendoscopés single-cell resolution to a conventional microscope, we imaged the same live neuron simultaneously with the microendoscope and an inverted microscope (IX 83, Olympus) in widefield epifluorescence mode. The epifluorescence microscope used a 490 nm LED light source (CoolLED, Andover, United Kingdom) and a 405/488/561/635 BrightLineLaser quad band filter set (F66-866, AHF). A cover slip with GFP positive neurons was placed onto the sample stage and imaged with a 40x objective (UPLXAPO40XO, Olympus, oil immersion, NA 1.4). Upon identifying a viable cell in the inverted microscope, this microscope was used to center the microendoscopés fiber above the same cell. Then images were taken with the epifluorescence microscope (exposure time 219 ms) and the microendoscope (2.5 s exposure time). For display [Fig. 3(c)], 5 images were averaged.

To characterize the resolution of the microendoscope we imaged 1 µm yellow-green (505/515) FluoSpheres (Life Technologies Corporation, Eugene, USA) suspended on poly-D-lysine (Sigma-Aldrich, St. Louis, Missouri, USA) coated microscopy slides [Fig. 3 (d)]. First a 100 µl droplet of 5 mg/ml poly-D-lysine in water was placed on a Superfrost plus adhesion microscope slide (Epredia, Essendonk, Netherlands) and left for 10 min at room temperature. The droplet was washed off with tap water and after drying a 10 µl droplet of a 1:5000 dilution of FluoSpheres in water was placed on the poly-D-lysine residue. After 10 min incubation at room temperature the slide was rinsed with tap water. The sample was not further mounted to allow for optimal access to the imaging surface with the microendoscope.

### 2.4 Spectra

Excitation spectra (Supplementary Fig. S1) were measured with a fiber-coupled spectrometer (CCS200, Thorlabs). Emission filter spectra were obtained from the vendor (AHF.de), the eGFP spectrum was obtained from fpbase.org [35, 36].

### 2.5 Animals

We used two mouse models to validate the microendoscope’s capabilities for in-situ imaging of cellular signals. These mice expressed cell specific fluorescent biosensors in either the kidney or the trachea. Additionally, a third mouse model, ubiquitously expressing the ratiometric sensor mito-roGFP2-Orp1, was used to confirm the multichannel capability of the system in whole explanted organs.

All animals that were part of our mouse models were bred and kept in facilities approved by the Saarland State Office for Health and Consumer Protection. Mice were kept in a 12 h light/12 h dark cycle with ad libitum access to water and food. It was an open colony and mice were kept in groups of 2 to 5 animals per cage. Male and female mice were used for the measurements. Handling and care of all animals was conducted in accordance with the German guidelines for the care and use of laboratory animals. All animals were killed according to our permit for euthanization for organ collection for scientific purposes (§ 4, paragraph 3 of the German animal welfare act). All experiments were performed on post-mortem mice immediately after death.

### 2.6 Podocyte calcium imaging

We recorded functional single-cell data in the kidney from mice that expressed the calcium sensor GCaMP3 in NPHS2 positive cells, i.e. podocytes. These cells were targeted by mating a NPHS2-Cre mouse [37] (Jackson strain number 008205) with a mouse carrying transcriptionally silenced GCaMP3 gene in the ROSA26 locus [38] (Jackson strain number 028764 kindly provided by Dr. Dwight Bergles, Johns Hopkins University School of Medicine, USA). Cre-mediated recombination allows for exclusive expression of GCaMP3 in renal podocytes [Fig. 4(c)]. All podocytes recorded were heterozygous for the GCaMP3 allele.

**Fig 4.**
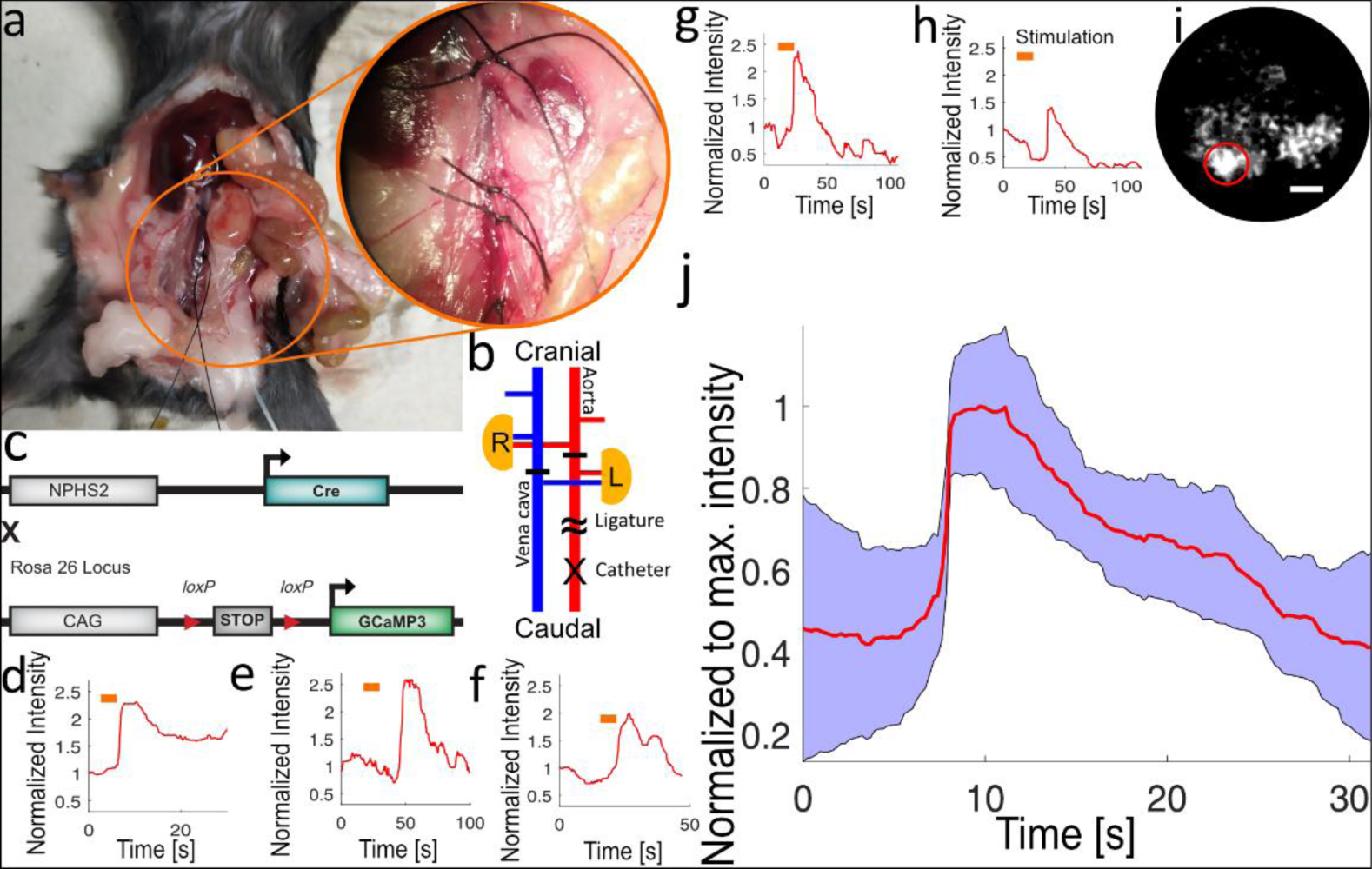
Functional calcium signals in mouse kidney. a) Surgical set-up and b) schematic view of the surgical procedure. The surgery aims to inject stimulation solution directly into the kidney to reduce signal latency. Solution injected into the infrarenal aorta through the catheter (X in b) can only flow through the kidney because all other vessels are ligated (**–** in b). The catheter is held in place by two ligatures (∼ in b). The microendoscope was placed onto the kidney and, after identifying individual podocytes by their basal GCaMP3 fluorescence, the kidney was flushed with 250 µl of 10 µM Angiotensin 2. c) Genetic background of mice. d–h) Individual calcium responses to Angiotensin 2 stimulation showed an abrupt signal increase, followed by a plateau phase before the calcium decrease (d–f) or an immediate calcium decrease (g, h). i) Processed image recorded by the microendoscope corresponding to trace h. Scale bar 20 µm. j) Average signal (red) of 5 stimulations, one per animal, with standard deviation (blue region). All traces were recorded at 4.3 frames per second.

To provide direct injection access to the kidney for stimulation of the podocytes, the abdominal cavity was opened after cervical dislocation. An arterial catheter (Intramedic PE10, BD laboratories, Mississauga, Canada) was placed in the infrarenal aorta between the renal and inferior mesenteric arteries. Ligatures were placed around the superior mesenteric aorta and the inferior vena cava, cranial to the renal vein. The renal capsule was removed. This method was adapted from Czogalla et al. [39] and ensured that the reagent perfused the kidney directly and reached the podocytes through the microvasculature. All surgical material was purchased from Fine Science Tools (Heidelberg, Germany). To image podocytes, fat and connective tissue surrounding the kidney were pushed aside and the microendoscope probe was suspended above the kidney with the micromanipulator. To locate GCaMP3 positive podocytes, the probe was slowly lowered toward the kidneýs surface until a signal was detected on the camera. The time from animal sacrifice to imaging was approximately 60–80 minutes, which is similar to established methods of kidney transplantation, indicating that the kidney retains viability [40].

For podocyte stimulation the intact kidney was perfused through the arterial catheter with 250 µl of 10 µM human angiotensin 2 (Sigma-Aldrich, Darmstadt, Germany, A9525), dissolved in custom kidney buffer (10 mM NaHCO_3_, 15 mM KCl, 100 mM KH_2_PO_4_, 198 mM D-glucose, adjusted to pH 7.0 with KOH). Before the first stimulation and between stimulations the kidneys were flushed with 250 µl of kidney buffer to flush autofluorescent blood out of the kidney and to ensure that the stimulant from the previous stimulation would not interfere with the current stimulation.

All time series were acquired with 200 ms exposure time (4.3 frames per second), using the 475/28 nm excitation light source and the 511/20 emission filter. The power of the 475/28 nm LED was 47 ± 3 µW in the imaging plane.

### 2.7 Brush cell calcium imaging

In the mouse trachea we recorded the response of single TRPM5-positive cells, i.e. brush cells, to the bitter substance denatonium benzoate. Brush-cell-specific expression of the calcium sensor GCaMP3 was achieved by mating a TRPM5-Cre mouse [41] with a mouse carrying a transcriptionally silenced GCaMP3 gene in the ROSA26 locus [38] (Jackson strain number 028764) [Fig. 5(b)]. Cre-mediated recombination results in GCaMP3 expression exclusively in TRPM5 expressing cells. All brush cells imaged were heterozygous for the GCaMP3 allele.

**Figure 5.**
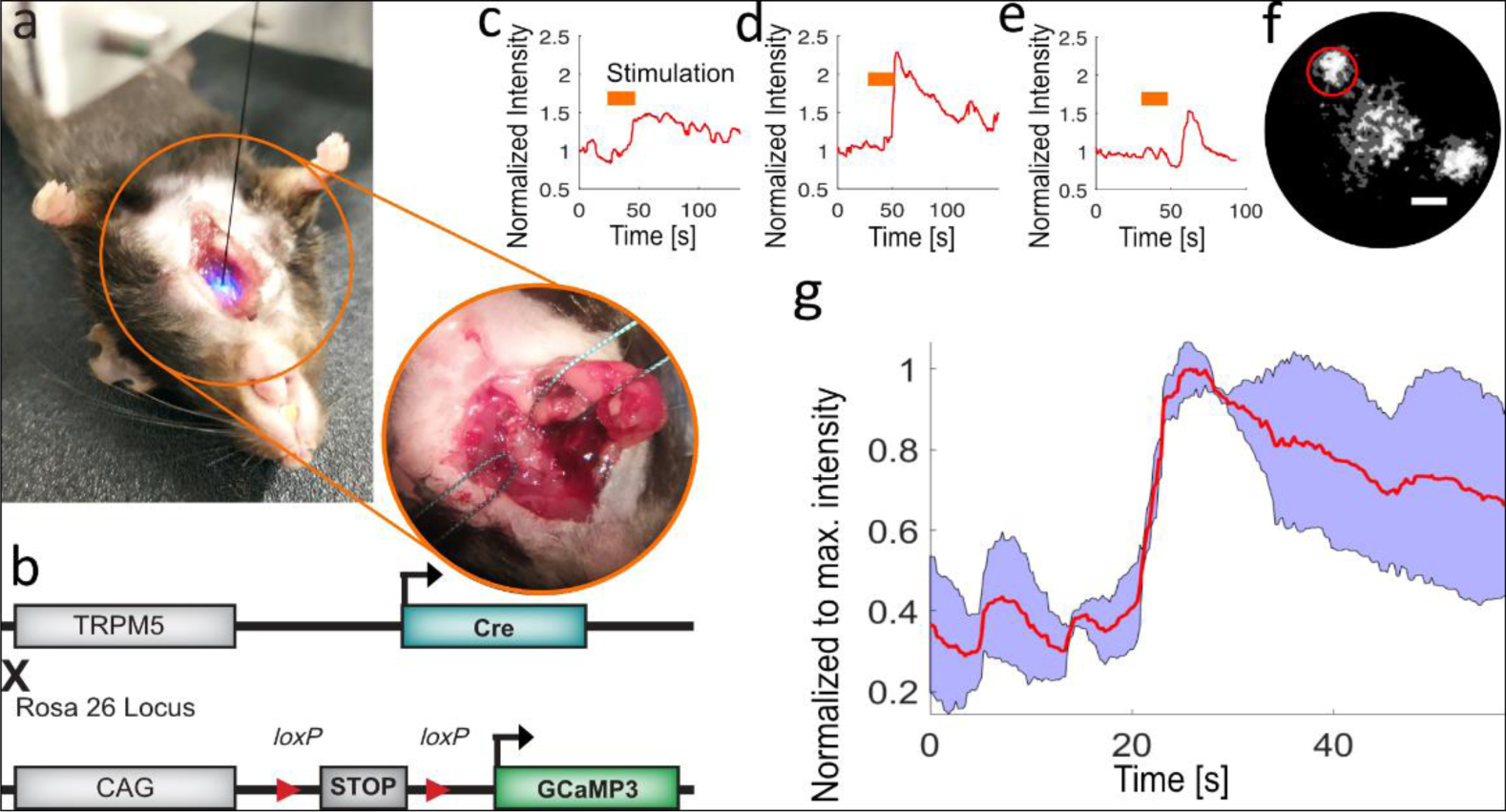
Functional calcium signals in mouse trachea. a) After insertion of the microendoscope probe into the trachea, the brush cells were excited with 4 µl of 10 mM denatonium. b) Genetic background of the mice. c–e) Individual calcium responses to denatonium stimulation showed an abrupt signal increase, followed by a long plateau (c) or an immediate signal decrease (d, e). f) Processed still image of video recorded by the microendoscope corresponding to trace e. Scale bar 20 µm (see Supplementary Movie 1). g) Average signal (red) of 3 stimulations, one per animal, with standard deviation (blue region). All traces were recorded at 4.3 frames per second.

We imaged functional signals of brush cells in the mouse trachea with the microendoscope through a small ventral cut that was prepared as follows: Animals were killed by cervical dislocation after which the throat was shaved. Then, the head was stretched with a piece of string, which was placed behind the animal’s incisors, pulled taught, and taped. Scissors were used to cut through the skin, revealing the salivary glands. The salivary glands were carefully pushed aside with blunt forceps and a pair of Dumont forceps was used to pierce the sternothyroid muscles. After pushing aside the opened muscle the trachea was cut open ventrally and a piece of single filament string was tied around a single cartilage ring on either side of the cut to gently pull the trachea further open. This filament was then taped down to keep the trachea continually open for a large accessible region [Fig. 5(a)]. The surgery was performed using a surgical microscope (Swivel arm, Zeiss, Jena, Germany). Experiments were performed within 20 minutes of death to maintain brush cell viability.

During imaging, the brush cells were stimulated with 4 µl of denatonium benzoate diluted to 10 mM in Tyrode buffer (130 mM NaCl, 10 mM HEPES, 10 mM D-glucose, 5 mM KCl, 1 mM MgCl2, 8 mM CaCl2, 10 mM sodium pyruvate, 5 mM NaHCO3) placed with a pipette onto the tracheal lumen.

All time-series were acquired with 200 ms exposure time (4.3 frames per second), using the 475/28 nm excitation light source and the 511/20 emission filter. The power of the 475/28 nm LED was 47 ± 3 µW in the imaging plane.

### 2.8 Brush cell distribution

To determine the brush cell distribution in the trachea, confocal images were acquired of a whole-mount trachea from a mouse expressing ChAT(BAC)-eGFP in TRPM5 positive cells (Supplementary Fig. S3). A similar breeding scheme as above was used: another mouse exhibiting the same TRPM5-Cre system as above was crossed with a mouse that carried a silenced GFP gene (Jackson strain 007902) instead of a silenced GCaMP3 gene.

To evaluate the spacing a whole-mount trachea sample for confocal imaging was prepared as described in the 2019 work of Hollenhorst et al. [21]. Brush cells were imaged by excising a whole mouse trachea from an animal with Chat-eGFP positive brush cells, which was then cut into three sections along the grain of the cartilage rings. The images (Supplementary Fig. S3) were taken using the 488 nm laser and 525/50 emission filter of the confocal/STED microscope described in the work of Staudt et al. [42].

### 2.9 Neuronal cell culture

To test the microendoscopes multichannel capabilities in cell culture using a static fluorophore (mKate) as a motion correction reference in addition to a calcium sensor (GCaMP6f), primary neuronal cell cultures were prepared as described previously [43]. Briefly, hippocampi were dissected from brains of newborn pups (P0) and digested for 20 min at 37 °C with 10 units of papain (Worthington, USA), followed by gentle mechanical trituration. Neurons were seeded at a density of 300 cells/mm^2^ onto 25 mm cover slips coated with 0.5 mg/ml poly-D-lysine (Sigma-Aldrich). Cultures were maintained at 37 °C in an incubator, humidified, with 95% air and 5% CO_2_ in NBA (Invitrogen, ThermoFisher Scientific, Waltham, Massachusetts, USA), supplemented with 2% B–27 (Sigma-Aldrich), 1% Glutamax (Invitrogen), and 1% penicillin/streptomycin (Invitrogen). Neurons were transfected with adenovirus after 5 days in culture and recorded between the 12^th^ and 16^th^ day in culture.

Neurons were imaged in extracellular solution (ECS, 130 mM NaCl, 10 mM NaHCO_3_, 2.4 mM KCl, 2 mM CaCl_2_, 2 mM MgCl_2_, 10 mM HEPES, 10 mM D-glucose, adjusted to pH 7.3 with NaOH). To evoke membrane depolarizations, cells were stimulated by perfusing the culture with high KCl ECS (90 mM NaCl, 10 mM NaHCO_3_, 80 mM KCl, 2 mM CaCl_2_, 2 mM MgCl_2_, 10 mM HEPES, 10 mM D-glucose, adjusted to pH 7.3 with NaOH).

Time-series of these neuronal cultures expressing GCaMP6f and mKate2 [Fig. 6(a)] were recorded using the 435/15, 520/10, 595/15, 695/30 quad band filter. Samples were illuminated alternatingly with 475/28 nm (GCaMP6f, odd frames) and 555/28 nm (mKate2, even frames) with an exposure time of 200 ms each. These frame-interleaved two-channel time series were acquired at a raw frame rate of 4.3 frames per second, which results in an effective frame rate of 2.15 frames per second per channel. The power of the 475/28 nm LED used for GCaMP excitation was 47 ± 3 µW in the imaging plane and for the 555/28 LED used for mKate2 measurements 54 ± 1.9 µW.

**Figure 6.**
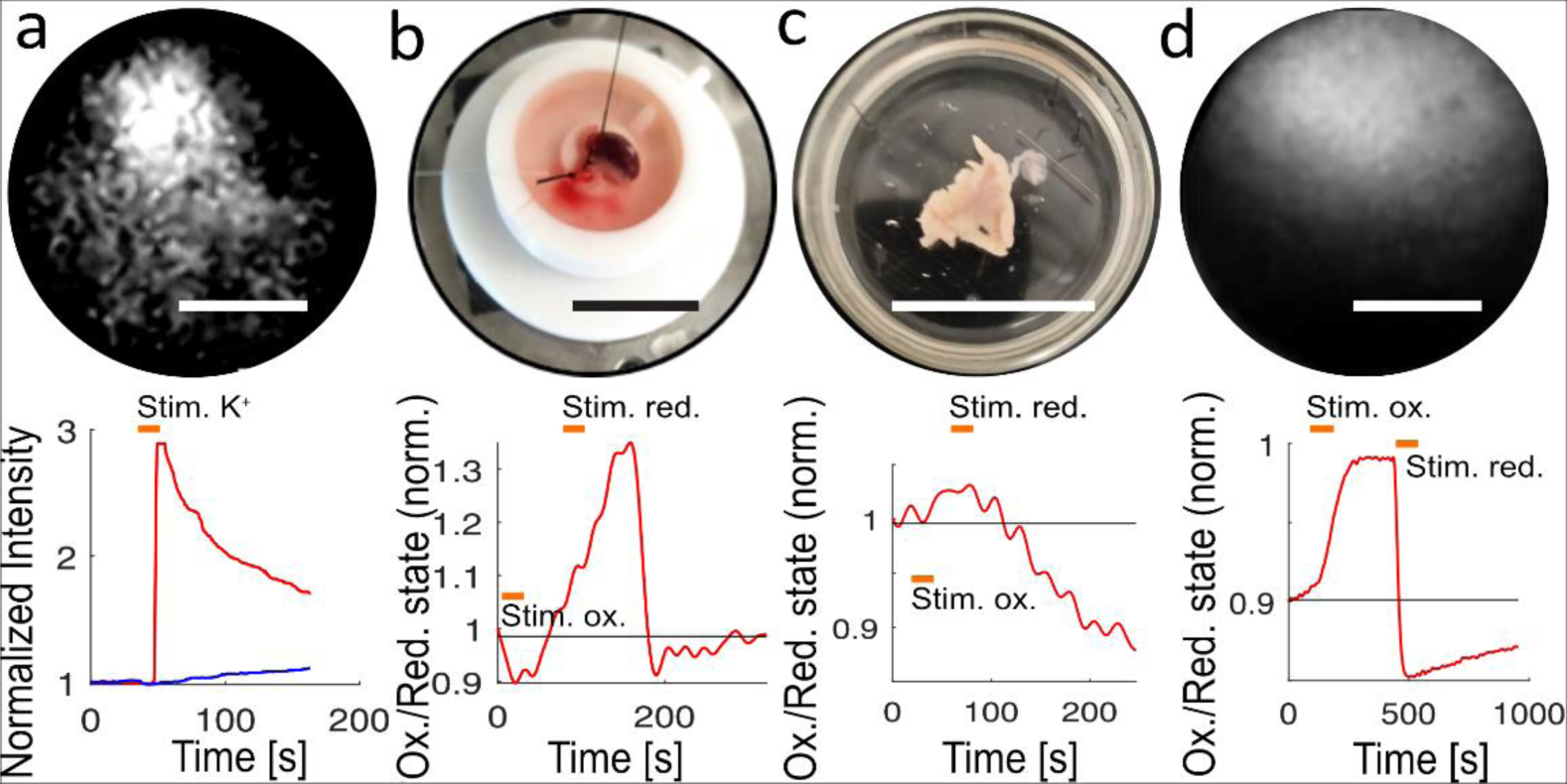
Multichannel recordings. a) Upper panel: Single neuronal soma after activation seen through the microendoscope. Lower panel: Calcium Response (red) of a single neuron to 80 µM KCl, while the frame-interleaved reference (blue) remains constant as expected. b) Upper panel: Excised kidney. Lower panel: Frame interleaved ratiometric recording of redox state changes in an excised kidney upon stimulation with oxidizing agent (Diamide, final conc. 2 mM) followed by reducing agent (Dithiothreitol final conc. 10 mM). c) Upper panel: Excised pancreas. Lower panel: Ratiometric recording of roGFP2-Orp1 redox state changes in an excised pancreas upon stimulation with oxidizing agent (Diamide, final conc. 2 mM) followed by reducing agent (Dithiothreitol final conc. 10 mM). d) Upper panel: Pancreatic islet seen through the microendoscope. Lower panel: roGFP2-Orp1 redox state changes are much faster and larger in the islet due to faster diffusion. Scale bars a, d 50 µm, b 5 mm, c 25 mm.

For optical characterization of the microendoscope the neuronal cell culture used was prepared in the same manner as described above using the mouse line “TRPC5–τ-green fluorescent protein knockin line” described in the work by Schwarz et al. [43].

### 2.10 Ratiometric redox imaging in whole organs

Ratiometric imaging with the microendoscope was validated using transgenic mice expressing the genetically encoded ratiometric H2O2 sensor mito-roGFP2-Orp1 targeted to mitochondria. These mice were previously described in detail in the work of Fujikawa et al. [33].

We recorded ratiometric redox signals with the microendoscope from several excised organs prepared as follows: Animals were killed by cervical dislocation. The pancreas and kidney were excised by opening the abdominal cavity and removing the respective organs with surgical scissors. The animal’s abdomen was shaved, and the skin was grasped with a blunt forceps. Surgical scissors were used to cut the skin horizontally. Once opened, the skin was cut vertically to expose the abdominal muscles, which were likewise cut. Upon the exposure of the abdominal cavity the target organ was identified, and the surrounding connective tissue and fat were removed with blunt forceps. The organ was cleared to the point where the only points of attachment were its major vessels, which were then cut in a single stroke to remove the organ quickly. During the surgery the organs were kept moist with Krebs-Henseleit buffer (described below).

For all ratiometric imaging the excised organs or islets were suspended in Krebs-Henseleit buffer (KHB, 120 mM NaCl, 4.8 mM KCl, 1.2 mM MgCl_2_, 2.5 mM CaCl_2_, 24 mM NaHCO_3_, 5 mM HEPES, 10 mM glucose, 1 g/l BSA, pH adjusted to 7.35–7.40 with NaOH) and stimulated with tetramethylazodicarboxamide (“Diamide”, Sigma-Aldrich, D3648, final concentration 2 mM) in KHB and with DTT (Sigma-Aldrich, D9779, final concentration 10 mM) in KHB. For stimulation Diamide was added to the submersion buffer to reach the final concentration of 2 mM (50 µl of 200 mM Diamide for a final concentration of 2 mM in 5 ml). For subsequent stimulation with DTT the buffer was exchanged to record a baseline between stimulations and DTT was added to reach the final concentration of 10 mM (100 µl of 500 mM DTT for a final concentration of 10 mM in 5 ml).

For the ratiometric mito-roGFP2-Orp1 imaging [Fig. 6(b–d)] we used the single-band filter. The samples were excited for 200 ms with the 395/35 nm source in odd frames and for 200 ms with the 475/28 nm source in even frames. The power of the 475/28 nm LED was 47 ± 3 µW in the imaging plane. For the 395/25 LED the power was 14 ± 1 µW.

Images were acquired at 1.8 frames per second per channel. In this case the reduction in frame rate compared to single-channel recordings was slightly greater than the obvious factor two, due to overhead for live display of both channels for visual control.

### 2.11 Ratiometric redox imaging in pancreatic islet culture

To test the system’s capabilities for ratiometric imaging outside the whole organ, we used a pancreatic islet culture. To prepare the culture, the abdomen of the mouse was fully opened after cervical dislocation, and the ampulla was pinched off. For perfusion of the pancreas, a small incision was made in the common bile duct and a collagenase solution [0.63 mg/ml collagenase P (Ref: 11213865001), Roche], dissolved in KHB [44] supplemented with 1% (vol/vol) penicillin/streptomycin was injected with a syringe. The inflated pancreas was excised and further digested in additional collagenase solution for 20 min at 37 °C; this temperature is maintained in a water bath in which the tube containing the pancreas-collagenase solution is placed. After mechanical homogenization of the pancreatic tissue by shaking, it was washed three times with KHB. Pancreatic islets were purified from exocrine tissue by hand-picking and cultured in RPMI 1640 (Ref: 21875034, Gibco^TM^, ThermoFischer Scientific) supplemented with 10% fetal bovine serum (Ref: 10270106, Gibco^TM^, ThermoFischer Scientific) and 1% (vol/vol) penicillin/streptomycin (Ref. P4333; Sigma-Aldrich) at 37°C and 5% CO_2_ overnight before measurement.

After overnight incubation, islets were placed in groups of 10 on a poly-L-lysine (Sigma, P4707) coated petri dish filled with KHB. The microendoscope was placed on top of an individual islet while the baseline was recorded. Subsequently, Diamide (final concentration 2 mM), followed by DTT (final concentration 10 mM) were then added to the solution, and the kinetic behavior was recorded with the microendoscope.

Imaging parameters were the same as for the whole pancreas.

## 3. Results

### 3.1 Optical Characterization of the Microendoscope

In a first step, we verified the specifications of the microendoscope experimentally. The theoretical magnification of the whole system is expected to be 55.56, based on the specifications of the objective (f=3.6 mm) and tube lens (f=200 mm) that image the fiber end to the camera [Fig. 1(a, b)] and of the GRIN lens with a nominal magnification of one [Fig. 1(c, d)]. The measured magnification (sample to camera) was 55.44 ± 0.53. The pixel size corresponds to 0.2883 ± 0.0029 µm per side in the sample plane. We found a diameter of 147.5 µm ± 1.7 µm for the FOV.

The imaging diameter of the fiber, where the individual cores are housed, should be 145 µm (corresponding area 16,500 µm²) and should contain 1600 cores with a tolerance of ± 10% according to the manufacturer. We counted the cores, which appear as individual bright spots under homogeneous illumination and found 1460 cores, thus one core per 11.3 µm². The center-to-center distance is accordingly expected to be 3.61 µm, if the cores are arranged in a perfect hexagonal lattice. Experimentally, we found an average distance of 3.71 ± 0.08 µm [Fig. 3(a)] and a core diameter of 2.34 ± 0.06 µm.

To validate that the microendoscope resolves single cells, we simultaneously imaged the same neuron from above through the microendoscope and for comparison from below through an inverted epifluorescence microscope. Not only the soma, but even the neurites are resolved [Fig. 3(c)].

Although resolving cells and subcellular structures is the final goal for the microendoscope, a precise characterization of the resolution is only possible with test samples such as fluorescent beads. The optical resolution of the microendoscope should theoretically be 592 nm for a wavelength of 485 nm, given the numerical aperture of the GRIN lens of 0.5 specified by the manufacturer (0.61*485 nm/0.5). Due to the much larger spacing of the fiber cores of 3.71 ± 0.08µm [Fig 3(a)], the effective resolution of the system is limited by the sampling of the individual fibers.

To characterize the imaging capabilities of the microendoscope experimentally, we imaged fluorescent beads with a diameter of 1 µm. These beads appear only on a single fiber core [Fig 3(d.1)]. Moving the fiber by one core-distance lets the bead appear on the neighboring fiber core [Fig 3(d.2)]. After the image processing that removes the individual cores, the bead appeared with a width of 4.67 µm (measured as full width at half maximum, FWHM), Fig. 3(d.3). The average FWHM (5 beads) was 4.64 ± 0.05 µm; this is the effective resolution of the system.

### 3.2 Imaging calcium signals of podocytes in intact kidneys in-situ

After the optimization and characterization steps, we applied the microendoscope to image functional in-organ signals. In the first series of experiments, we recorded calcium responses of GCaMP3-expressing podocytes in the intact kidney. For hormonal stimulation of these cells, we inserted an arterial catheter into the infrarenal aorta of freshly euthanized mice. After occlusion of the superior mesenteric aorta, an injection through the catheter can only flow through the kidney, allowing for direct stimulation of the podocytes [Fig. 4(a–c)]. This was necessary because previous experiments with excised kidney (not shown) had revealed that the diffusion time of the stimulant (Angiotensin 2 as described below) into the podocytes was unpredictable and prolonged imaging (in the range of 30 min) resulted in photobleaching. We performed stimulations in five kidneys and analyzed these signals after image processing and filtering, the results of which are illustrated in Fig. 4(d–h). A representative image of one experiment illustrating how the podocyte appeared through the microendoscope is shown in Fig. 4(i). Stimulation evoked significant increases in the GCaMP3 signals in single podocytes although the exact time course varied. We found an approximately twofold increase in cell-specific fluorescence intensity upon stimulation. After stimulation, the signal was often preceded by a short (<10 s) dip in intensity before rising rapidly over the span of 1–3 s. Some signals exhibited a short (<10 s) plateau before falling over the next 15–20 s [Fig. 4(d–f)], whereas other signals lacked this plateau [Fig. 4(g, h)]. The signals are reproducible as seen in the signal averages plotted with their standard deviation [Fig. 4(j)].

### 3.3 Imaging calcium signals of tracheal brush cells in-situ

To verify the applicability of our novel microendoscope system for in-situ imaging in very small cavities that are usually inaccessible to larger instruments, we recorded functional calcium signals from GCaMP3-expressing brush cells in the mouse trachea. After opening the skin, salivary glands and the sternothyroid muscles were positioned laterally and the trachea was opened ventrally. A single cartilage ring on either side of the trachea was then used as an anchor for a single-filament string to further open the trachea and to increase the accessible surface area [Fig. 5(a)]. This was necessary because previous experiments had revealed that the number of brush cells increases from the dorsal to the ventral part of the trachea [22]. With the tilting sample/animal stage the animal was tilted ∼15° along the anterior-posterior axis. This gave access to the side of the tracheal wall where more brush cells are expected. We then stimulated the trachea locally with 4 µl droplets of 10 mM denatonium benzoate. The individual brush cells in the trachea responded to this stimulation with a time-dependent increase in intracellular calcium concentration, as seen by the increase of the GCaMP3 [Fig. 5(b)] signal [Fig. 5(c–e)]. At the peak an approximately 1.6-fold increase in cell-specific intensity was observed. The signal intensity increased over the first 1–3 s, sometimes plateauing, before beginning to fall over the next 20–50 s. Signal averages indicated reproducible results [Fig. 5(g)]. The average of all frames showed the position of the individual brush-cells [Fig. 5(f)]; the low pre-stimulation intensity of the cells, the intensity increase upon stimulation and the post-stimulation reduction in fluorescent intensity is apparent in the video [Fig. 5(f), Visualization 1]. To further corroborate that the observed signals are likely from single cells, we imaged the brush-cell distribution in a fixed whole-mount trachea (Supplementary Fig. S3). The individual brush cells were sparsely distributed in the trachea, indicating that the dynamic calcium signals stem from single cells.

### 3.4 Dual channel recordings

In the following experiments we explored the ability of our system to perform dual-channel recordings by imaging neuronal cultures expressing GCaMP6f and mKate2. Neurons were imaged with frame interleaved excitation at 475 nm (for functional calcium/GCaMP6f signals) and 555 nm (for the mKate2 reference channel). Each channel was plotted separately to show that upon induction of membrane depolarization by stimulation with 80 mM KCl, the functional trace of the GCaMP6f fluorophore showed a 2.5-fold increase in cell-specific fluorescence intensity due to Ca^2+^ influx through voltage-gated calcium channels (VGCCs) while the non-functional reference fluorescence remained largely unchanged [Fig. 6(a)], excluding artificial fluorescence changes, e.g., due to motion artifacts.

Finally, to demonstrate the microendoscopés capability to detect ratiometric signals, we recorded redox signals with the fluorophore roGFP2-Orp1 in whole, excised organs, as well as cultured pancreatic islets. roGFP2-Orp1 is a modified form of GFP that is excitable at two wavelength ranges, with the effectiveness of either being determined by the oxidation status of the fluorophore specified by the presence of H_2_O_2_. In the presence of an oxidizing agent the excitation efficiency of roGFP2-Orp1 is high at shorter wavelengths (∼400 nm), whereas a reducing agent leads to a higher excitation efficiency at longer wavelengths (∼490 nm) [29].

For ratiometric measurements of organs, the pancreas and kidney of a ubiquitously mito-roGFP2-Orp1 expressing mouse were excised. They were washed in Krebs-Henseleit buffer (KHB) to remove superficial blood, which could otherwise stick to the lens and exhibit of autofluorescence [32]. All specimens were then imaged in KHB with frame-interleaved excitation at 395 nm and 475 nm. Oxidation of the samples was attained by the application of Diamide (final concentration 2 mM), and reduction was achieved with dithiothreitol (DTT, final concentration 10 mM). The resulting movie was split into two separate time series, sorted by the excitation color. For each time series, ROI-based intensity traces were generated (see Methods) and their ratio was calculated for each time point. This final trace was then normalized to the mean of the first 50 values.

The ratiometric trace exhibited the expected changes in the fluorescence characteristics of roGFP2-Orp1 in whole pancreas and kidney [Fig. 6(b, c)]. Notably, the organ recordings were time-delayed, likely due to the time required for the reagent to diffuse into the tissue. In comparison, the experiments performed in isolated pancreatic islets yielded rapid responses with minimal delay [Fig. 6(d)], as expected, based on shorter diffusion times of the oxidizing/reducing agent in culture.

## 4. Summary and Discussion

We have developed and characterized a microendoscope capable of imaging functional cellular fluorescence signals in cavities as small as a mouse trachea (Figs 1, 3) and programmed an appropriate data processing pipeline (Fig. 2). We validated the microendoscope in several organs using diverse biosensors (Fig. 4–6). Additionally, we exploited its multichannel capabilities for ratiometric and reference channel imaging in intact organs and in neuronal and pancreatic islet cultures, highlighting the flexibility of the system (Fig. 6). To investigate organs such as the kidney it is usually required to either explant and stabilize the kidney [26, 45] or to use techniques that cannot resolve single cells [39, 46–49], whereas the microendoscope enables flexible and cellular-resolution imaging of the kidney in-situ.

The capability of our microendoscope to acquire functional single-cell data was particularly relevant as it provides a system that is small enough to enter spaces such as mouse tracheae that are inaccessible for conventional microscopy or larger microendoscopes. It thus enabled us to quantify cellular responses in tracheal brush cells (Fig. 5), which previously had to be cultured or excised and incubated to investigate their functional capabilities [18, 22, 50]. Our microendoscope enabled their investigation in-situ and yielded fluorescence signals comparable, in shape and time course, to those obtained with a high-resolution microscope [18]. Additionally, we were able to visualize and record calcium signals in renal podocytes (Fig. 4). The calcium imaging of single cells shows that the microendoscope is capable of resolving these cellular signals and suggests that they may exhibit intercellular variability. We observed kidney podocyte signals with a longer signal peak (Fig. 4 d–f) compared to others [Fig. 4 (g, h)], as well as tracheal brush cell signals with a prolonged peak [Fig. 5(c) vs. (d, e)].

Podocytes are located superficially in the renal cortex, but still deeper than the superficial layer, in approximately the 2^nd^ to 5^th^ cell layer [51]. With a working distance of 110 µm the microendoscope is not only suitable for imaging superficial cells but can also record from deeper layers. The microendoscope excites via a LED source in a widefield epifluorescence configuration. It therefore lacks the optical sectioning capabilities of conventional confocal or two-photon microscopes. However, a confocal or two-photon microscope requires the sample to be brought to it, whereas the microendoscope fiber can be maneuvered to the sample and can still detect individual cellular signals at some depth. Endoscopes using confocal and two-photon configurations have been conceived. The optical sectioning capability comes at the cost of much larger endoscope diameters, relying on more complex probe set-ups on the imaging end, and demonstrations of biological or even functional imaging are often missing [10, 13, 52, 53]. Whereas two-photon microscopes allow imaging at a depth of several hundred micrometers [54] the flexibility of the endoscope allows it to bring the instrument much closer to the cells of interest and to image a sample from unconventional angles. We demonstrated in the trachea that the microendoscope is capable of imaging the same cells as a two-photon microscope, with a working distance of only 110 µm.

Our microendoscope is tailored for functional imaging of cellular signals in the whole animal. Two features here are of particular relevance: First, the multi-channel capability. It allows ratiometric measurements and measurements with a reference channel to distinguish functional signals from motion artifacts, which are always a concern in in-vivo measurements [55]. Second, its small size, which makes deeper organs accessible with minimal lesions and allows imaging in small cavities. Combined with cellular resolution (Fig. 3) the small size of the microendoscope makes it uniquely suited for imaging small animals such as mice. This combination allows the system presented in this paper to play an important role in the field of intravital imaging.

Longitudinal studies of organs at the cellular level are made possible by the ability to reliably image organs that lie in soft tissue or are generally difficult to access. Currently, longitudinal studies rely on solid structures, such as the skull, to mount a camera system such as the Miniscope [56] or they rely on non-invasive techniques that lack cellular resolution [47, 49, 57]. Alternatively, a population of animals is imaged at different time points, but each animal only once [45]. Investigating for example the progression of kidney injury, kidney disease or the effect of heart disease on the renin-angiotensin-aldosterone-system at the cellular level could lead to new insights and disease models, while reducing the number of animals required for such studies. The microendoscope presented in this study is uniquely suited to image cellular responses in soft tissue environments lacking a hard anchor point, such as the skull. While bigger systems, fixed to the skull allow for cellular microscopic imaging with a larger field of view and better resolution [56] our microendoscope provides a solution for functional imaging with single cell resolution in diverse organs.

Additionally, the flexibility of our microendoscope lends itself to combination with other imaging techniques to perform simultaneous multi-organ in-vivo imaging. To characterize the microendoscope presented here, we imaged the same cell with a conventional microscope and with the microendoscope [Fig. 3(c)]. This approach can be extended to multi-site imaging in the same sample or animal. In such a set-up a larger, more complex optical system could be used to image (and eventually optogenetically stimulate) one organ, such as the heart, while the smaller, more flexible microendoscope is used to simultaneously record induced functional signals from a second organ (such as the kidney). Such flexibility will be an asset in advancing biological investigations around the organs established here, such as the kidney, trachea, pancreas, and beyond.

## Supplementary Material

Extended versions of the autofluorescence spectrum [Fig. 1(e)] overlaid with excitation spectra, raw calcium imaging data overlaid with the filtered trace [Fig. 4(e), Fig. 5(e)] and additional information on confocal imaging of tracheal brush cells to show their sparse distribution are presented in the supplementary material.

## Author Declarations

### Funding

TAD acknowledges a fellowship of the Studienstiftung des deutschen Volkes. MAL, GKC and PL acknowledge support by the NanoBioMed Seed Grant “In vivo fiber-based imaging” (Saarland University). GKC further acknowledges funding by DFG SFB TRR 152 project P22, as well as KR4338/1-2 and MIE acknowledges funding by Homfor2023 Nachwuchsförderung (Homburger Forschungsförderung, Saarland University).

## Acknowledgements

We would like to thank Emmanuel Ampofo for his kind support and valuable time, as well as Matthias W. Laschke for his support early in the project. Additionally, we thank David Stevens, Ahmad Lotfinia, Yasser Medlej and Jan Knobloch for their comments on the manuscript and figures. Finally, we would like to acknowledge Xin Hui for his support with the mouse lines and GRINTECH Gmbh (Jena, Germany) for their excellent technical support.

## Disclosures

The authors have no conflicts to disclose.

## Notes

### Competing Interest Statement

The authors have declared no competing interest.

