## Supplementary Figures for "Multicore-fiber microendoscopy for functional cellular in-organ imaging"

### Contents

Section S1: Additional filter overlay

Section S2: Raw traces of calcium signals

Section S3: Whole-mount trachea for density estimation of brush cells

### Section S1: Additional filter overlay

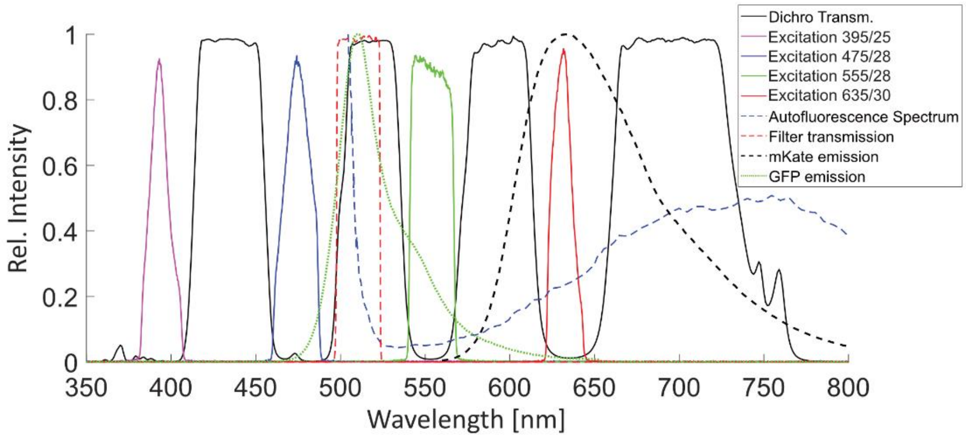


**Figure S1. Emission filter transmission overlaid with dichroic mirror transmission, autofluorescence spectrum and excitation wavelengths.** Single-band filter [cf. Fig. 1(e)].

### Section S2: Raw traces of calcium signals

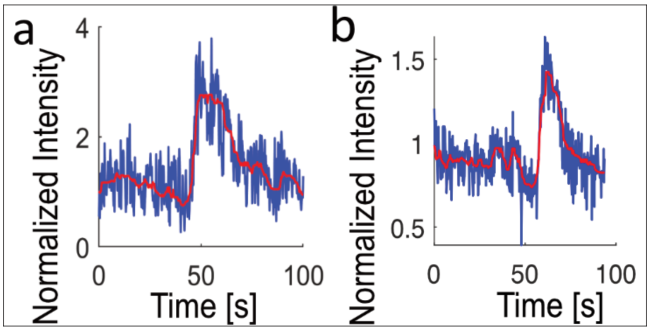


**Figure S2. Raw Data of calcium Imaging.** a) Single podocyte response to stimulation with angiotensin 2, blue trace shows the raw recorded calcium signal whereas the red trace shows the filtered signal [cf. Fig. 4 (e)]. b) Calcium response of a single tracheal brush cell upon stimulation with denatonium. Blue trace shows the raw recorded calcium signal whereas the red trace shows the filtered signal [cf. Fig. 5 (e)].

### Section S3: Whole-mount trachea for density estimation of brush cells

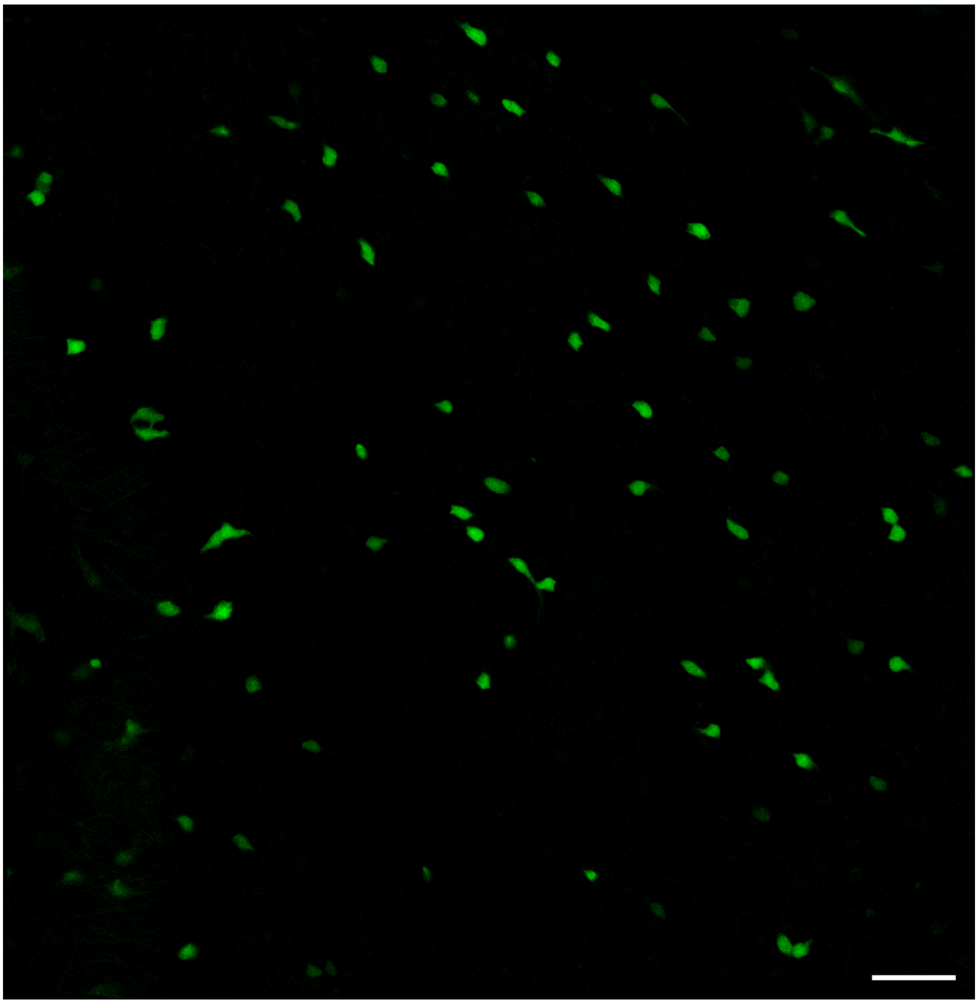

**Figure S3. Confocal image of brush cells in fixed whole-mount mouse trachea**, revealing a sparse distribution. Scale bar 50 µm.
